# Measuring *C. elegans* Ageing Through Non-Invasive Monitoring of Movement across Large Populations

**DOI:** 10.1101/2023.06.15.545090

**Authors:** Giulia Zavagno, Adelaide Raimundo, Andy Kirby, Christopher Saunter, David Weinkove

## Abstract

Finding new interventions that slow ageing and maintain human health is a huge challenge of our time. The nematode *Caenorhabditis elegans*, offers a rapid *in vivo* method to determine whether a compound extends its 2-3 week lifespan. However, the standard *C. elegans* lifespan assay is hard to scale for large screens. Lifespan analysis produces only one data point per animal with no information about health. Here we describe automated monitoring of movement from early to mid-adulthood as a healthspan-based alternative to measure ageing. Using our WormGazer™ technology, over 100 petri dishes containing *C. elegans* worms are imaged simultaneously and non-invasively by an array of cameras. This approach demonstrates that most functional decline in *C. elegans* occurs during the first week of adulthood. We find 7 days of imaging is sufficient to measure the dose-dependent efficacy of sulfamethoxazole to slow ageing, compared to 40 days required for a parallel lifespan experiment. Understanding any negative consequences of interventions that slow ageing is important. We show that the long-lived mutant *age-1(hx546)* stays active for longer than the wild type but it moves slower in early adulthood. Thus, continuous analysis of movement can rapidly identify interventions that slow ageing while simultaneously revealing any negative effects on health.

## Introduction

Ageing drug discovery faces specific and unique challenges. One challenge is that there are no fast cell-based systems to test whether a compound slows ageing. Instead, cell culture-based approaches depend on measuring aspects of cell biology thought to be important in ageing, known as the ‘hallmarks of ageing’ (López-Otín et al. 2013, López-Otín et al. 2023). This approach limits the targets to the bounds of current knowledge and to interventions that work on properties detectable in cultured cells. Ageing involves interactions of multiple biological systems at several levels (organs, tissues, cells, molecules), which can only be seen in whole organisms. Experiments in mice are time consuming, expensive, and constrained by ethical regulation. Thus, testing in a model organism such the nematode *Caenorhabditis elegans*, which ages in weeks, provides the opportunity to gain data from compounds in an *in vivo* ageing system. Another challenge in ageing drug discovery is that interventions would be given long-term to healthy people, and so must be completely safe. Any potential side effects should be detected early as possible in the drug discovery process.

At least 83% *C. elegans* proteins have human homologs (Lai et al. 2000), and its short lifespan allows easier study of ageingand how a drug behaves in an aged organism. Genetic analysis in *C. elegans* was used to discover that the Insulin/IGF-1 signalling (IIS) pathway modulates ageing across several species (Lin et al. 1997, Harrison et al. 2009, Partridge et al. 2011), showing that at least one mechanism of ageing is well conserved through evolutionary distance. *C. elegans* has also been used to show that many long-lived mutants experience physiological trade-offs that include reduced fecundity or delayed development (Walker and Lithgow 2003, Jenkins et al. 2004, Bansal et al. 2015, Zhang et al. 2016, Podishivalova et al. 2017). These findings highlight that measuring lifespan alone is not sufficient to understand whether an intervention will be beneficial in humans. We also need to be aware of any negative consequence on health.

The manual *C. elegans* lifespan assay is accessible and easily transferrable between research groups but can be subjective and difficult to standardise (Lucanic et al. 2017). The assay collects binary data on whether the worm is alive or not by checking for signs of movement. If a worm is not moving, then it is prodded with a platinum wire to look for a response. If there is no movement response, the animal is scored as dead. Technologies that enable automated image analysis can increase the amount of data collected in the same experiment while making more objective measurements and saving labour. These technologies assess time of death by measuring movement on solid media (Stroustrup et al. 2013, Churgin et al. 2017) or in microfluidic environments (Hulme et al. 2010, Rahman et al. 2020).

There are technologies (including those referred to above) that can provide parameters such as fecundity (Li et al. 2015), body bends (Cronin et al. 2005), and pharyngeal pumping (Rodriguez-Palero et al. 2018), which informs how health changes with age, providing a measure of healthspan. However, these technologies are either invasive, because animals have to be moved to a device for testing, or require animals to be kept in microfluidic devices throughout their adult lives and thus be in a liquid environment, which is known to produce different physiology to animals kept in a solid environment (Lev et al. 2019).

Here, we describe how health can be monitored non-invasively, on agar-filled petri dishes with live bacterial lawns, which are the conditions for standard lab culture. This technology, which we have named the WormGazer™, monitors several petri dishes simultaneously without moving the dishes or the cameras. By monitoring the worms from the L4 stage (1 day before adulthood) and with an experimental runtime of 7-10 days into adulthood, we show that measuring movement in this timeframe can detect improvements in health created by drugs that slow ageing. We further use the drug sulfamethoxazole (SMX) to show that delays to movement decline caused are clear in the first 7 days of imaging, whereas effects on survival in a concurrent parallel manual experiment take several times longer. Finally, we test the reported lifespan-extending compound alpha-ketoglutarate. This approach can be used in drug development to improve understanding of the effect of a compound in a whole organism, allowing more informed decisions about lead candidates and thereby accelerating progress.

## Results

### Monitoring the Movement of Large Populations of Worms Over Time

Multiple technologies have been designed to monitor worm movement but most are hard to scale because they either involve a single fixed high-power camera that worms need to be moved under in sequence, or stationary petri dishes, multi-well plates, or microfluidic devices with worms where a camera moves between them. These technologies present the challenges of precisely and consistently aligning moving cameras. To monitor a large number of worms simultaneously, we use a large array of low-grade cameras each connected to a single board computer rather than a small number of high-quality cameras. This distributed computing approach with no moving parts is designed to be robust and scalable. Using a series of sequential images to detect movement, this system can identify worms that move during the imaging window. It is not necessary to track each individual worm because measuring changes in behaviour at a population level is sufficient to measure ageing.

Each 6 cm petri dish of solid media is illuminated from above and imaged by a camera below it (Figure 1a). This camera is controlled by a connected single board computer. Every 5 minutes, each camera records 200 images in a 160 second time frame. The contrast between the worm and its translucent medium allows for the measurement of changes in pixel values (Figure 1b left). When the images from an imaging window are averaged (Fig. 1b middle left), subtracted from each frame, and then overlaid, a high contrast image is created representing the movement in the imaging window, where pixels which changed in value during the imaging window due to movement are denoted as white and the unmoving background denoted in black (Figure 1b middle right). Where and how far a worm moves during the imaging window is therefore determined by following the path of these bright pixels at their centre of mass (Weinkove et al. 2021). A threshold speed for detection is set to 10 μm s^-1^ to limit noise from other changes to the plate which the camera can pick up, such as worm trails in the bacterial lawn. There are some false positives – objects detected that are not worms, occasionally resulting in an apparent more than 100% worms moving at a single time point but these anomalies are not at a sufficient level to mask movement at a population level. All objects moving above the threshold are used to measure fraction of worms moving (number of objects), mean speed of moving worms (length of objects), and position of the worm on the plate (coordinates of object) (Figure 1b, right). There is an area limit for worm detection, denoted by the red ring (Figure 1b, left and middle right). This system allows for the non-invasive monitoring of worms in their normal laboratory setting.

**Figure 1.**
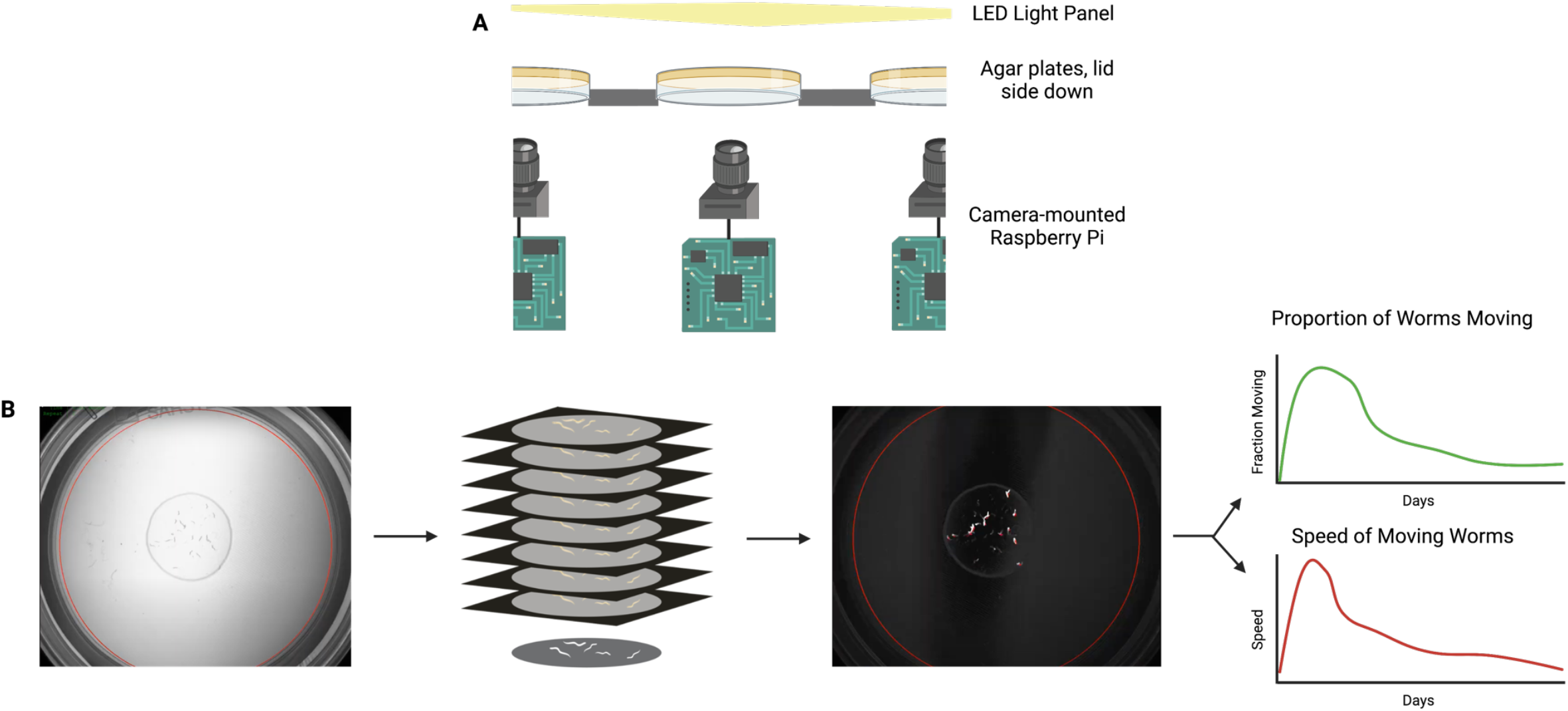
Schematic of the WormGazer™. (A) Petri dishes are placed in a temperature-controlled box with one imaging station per petri dish. This diagram represents how the camera is positioned to take images and is connected to a single board computer. The camera takes an image every 0.8 secs for 160 seconds and repeats this process every 5 minutes. The first level of image processing occurs on the single board computer. (B) A transmission image from the camera of a plate with 30 L4 worms (left). The images’ pixel values are averaged across 200 images in each imaging window (middle left), producing a high contrast image where movement can be detected (middle right). The fraction of animals moving and speed which the moving objects move are two main outputs (right). Created in Biorender.com.

As temperature is an important variable for ageing in *C. elegans*, and since changes in temperature can affect the moisture content of the petri dishes, the chamber containing the dishes is carefully temperature controlled using a feedback system of cooling by blowing externally chilled air to counteract the heat emitted by the lights and electronics. Extra heat is provided by small resistance heaters when necessary. Illumination is provided by white LED panels and are always on for imaging purposes. Data analysis occurs on locally connected servers, and Python scripts are used to create initial data outputs.

Once an experimental run is started, the boxes are left undisturbed for the duration of the run, which is between 7 and 14 days. The worms are placed on the dishes at the L4 stage, and the box normally runs at 24.0 °C. Under these conditions wild type worms become adults by 24 hours, show peak movement around 48 hours, and then begin their ageing-associated decline in movement and speed.

### Characterisation of the Movement of the Long-Lived *age-1(hx546)* Mutant

To test whether the system could detect changes due to ageing in *C. elegans* mutants established to show effects on lifespan, we compared the movement of the long-lived *age-1(hx546)* and the short-lived *daf-16(mu86)* mutants with a wild type (WT) control. A minimum of 298 worms across 10 dishes per condition were compared over a 12-day period. 2 μM FUdR was added to the media to prevent eggs from hatching.

The fraction of worms moving at any time is defined as what proportion of the population are moving during the 160s imaging window. The lines of the graph are smoothed, with shading of the standard error on the mean (SEM) (Figure 2a). Movement of the worms reaches a peak around day 2 so this timepoint is considered to be the start of age-related decline and is used as the starting point for the area under the curve (AUC) calculation, which produces the average time the worms spent moving in that time period.

**Figure 2.**
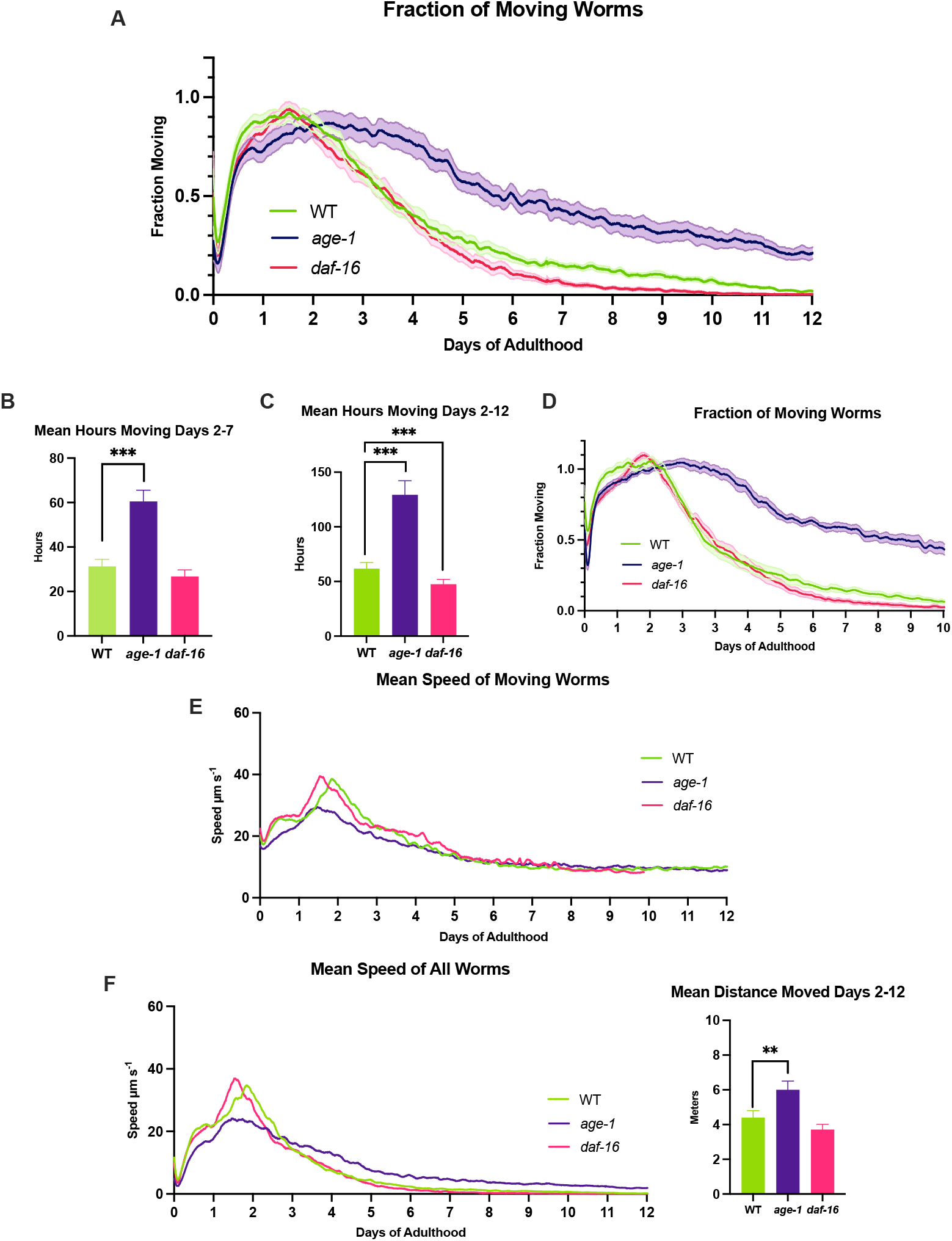
Movement Analysis of age-1, daf-16 and WT worms over 12 days. (A) The fraction moving graph shows the proportion of worms moving during the imaging window with SEM shading. n ≥ 298 worms, 10-12 petri dishes per condition. The area under the curve integration for day 2-7. (C) The area under the curve integration for days 2-12. (D) Fraction moving of an independent repeat, with SEM shading. n ≥ 420 worms, 14 petri dishes per condition. (E) The mean speed of moving worms. (F) The mean speed of all worms, which is a function of A and E. Plates with 2 μM FUdR on DM. *** = p <0.002, one-tailed test. Plotted in GraphPad.

The *age-1* mutant worms spend 93.5% more time moving between days 2 and 7 compared to the WT (Figure 2b). However, when the AUC is calculated for days 0.5-2, *age-1* mutants show a 12.1% lower fraction moving compared to WT (p < 0.05, Supplementary Table S2). This unexpected result suggests that the *age-1* mutant has fewer moving worms in early adulthood compared to WT.

Interestingly, the *daf-16* mutant showed no significant difference in average time spent moving in the first 7 days when compared to the WT, but when the AUC is calculated for days 2 to 12, it spent 23.2% less time spent moving compared to WT (Figure 2c). Overall, these results are consistent with slower ageing in the *age-1* mutant because it stays active for longer than the WT, while *daf-16* experiences a faster decline than the WT. The data show excellent reproducibility (Figure 2d).

The mean speed of moving worms is a function of how far each detected worm moves during the imaging window (Figure 2e) and does not consider those worms which do not move during the window. In this analysis, the *age-1* mutant had a 27% lower speed than WT at day 2 (p < 0.002, Supplementary Table S2), as well as reaching its maximum mean speed at day 1.53 compared to WT at day 1.94 (Figure 2e). Meanwhile, there was no significant difference in speed between WT and *daf-16*. After day 10, no more speed data are available for *daf-16* as at least one entire dish of worms had zero worms that moved above the detection threshold of 10 μm s^-1^.

When the fraction moving and mean speed of moving worms parameters are multiplied, it produces the mean speed of all worms (Figure 2f). This graph summarises the findings from this experiment, where *age-1* has a lower peak in speed but continues moving at higher and faster levels for longer, and that *daf-16* has an earlier and steeper decline than WT. The area under this curve represents the average distance the worms move and can be used for statistical analysis. A full comparison of the two repeats can be found in Supplementary Figure S1.

### Automated Movement Analysis vs Manual Lifespan

To compare movement analysis between the WormGazer™ and a manual lifespan assay, both approaches were performed in parallel. 26 dishes of 30 worms per condition were prepared with 50% analysed on the WormGazer™, and 50% by manual lifespan methods (n=390 set up per condition, per method). Worms were transferred to fresh dishes on days 7 and 14 with both methods. SMX was used as a positive control. SMX extends *C. elegans* lifespan in dose-dependent manner up to at least 256 μg/ml by inhibiting folate synthesis in OP50 *E. coli*. Inhibiting bacterial folate synthesis inhibits an *E. coli* activity that that accelerate ageing (Virk et al. 2016).

The manual lifespan experiment (Figure 3a) showed a statistically significant increase in survival with increasing SMX concentration (p < 0.0001 for all conditions, Wilcoxon test). Mean survival for 1 μg/mL SMX was 20 days, while 4 μg/mL and 8 μg/mL were 21 days compared to control which was 18 days (Supplementary Table S1). This experiment took 40 days to complete.

**Figure 3.**
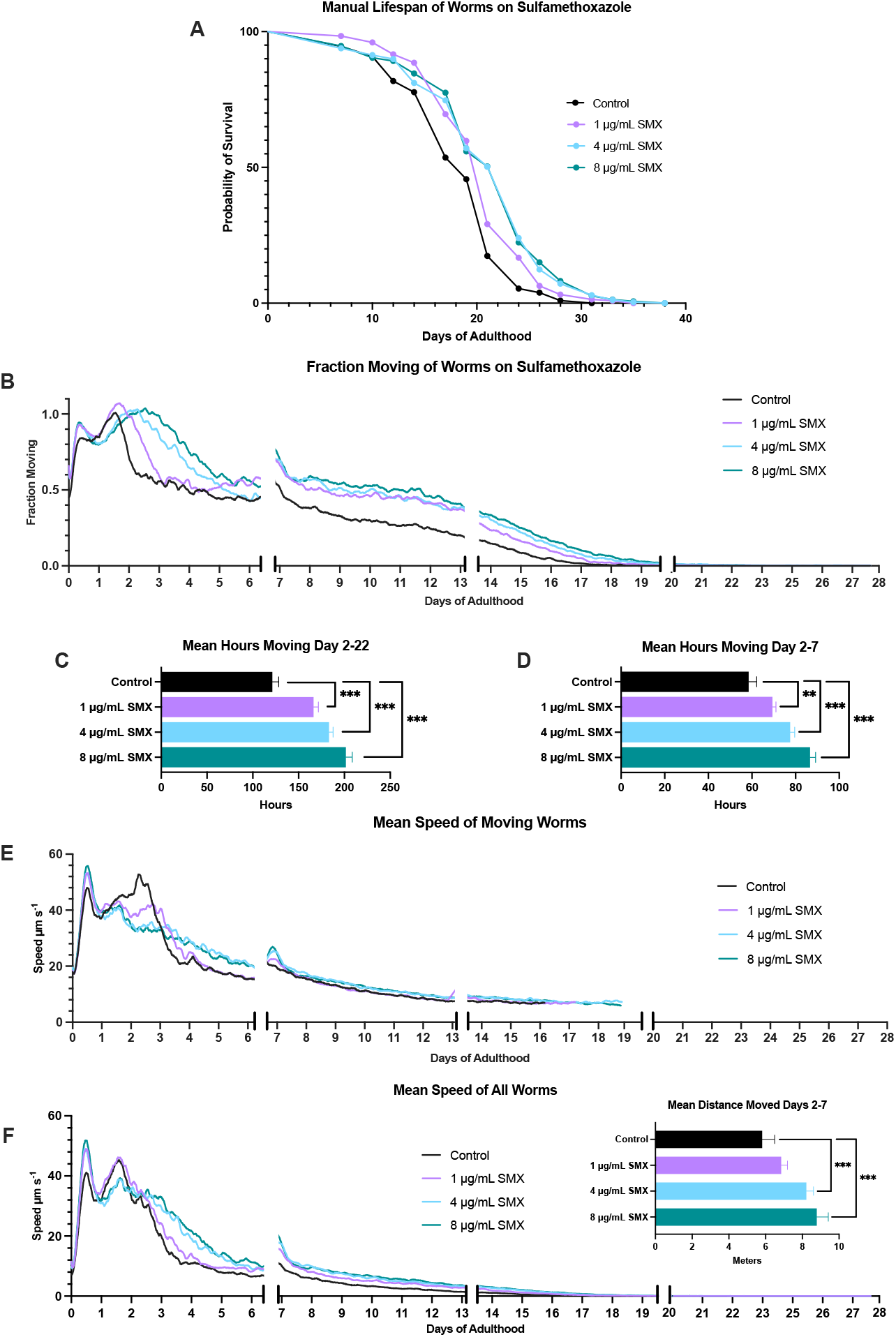
Manual lifespan versus WormGazer™ Movement Analysis. glp-4 sterile worms placed on as L4s, and were transferred on days 7 and 14, being placed either into the WormGazer™ or into the 24 °C incubator. (A) Survival curve of the manual lifespan method. (B) Fraction moving of the worms on the WormGazer™. An interrupted axis has been added to the fraction moving plot to indicate when the worms were transferred in the first two instances, while the last gap indicates when the machine was restarted. All these instances created bumps in movement which were smoothened by omitting 46 imaging periods. (C) Integration of area under curve is shown for days 2-22, and (D) days 2-7. The mean speed of moving worms (E) and the mean speed of all worms (F) which multiplies B and E. Manual scoring occurred every other day on weekdays from day 7 onward. ** = p < 0.01, *** = p < 0.002, one-tailed test. Compound added to agar. n ≥ 260 worms, 12 petri dishes per condition, per technique. Plotted in GraphPad.

In the WormGazer™ assay, worms showed an improvement in movement levels on all SMX concentrations (Figure 3b) that was statistically significant from day 2 onwards (p < 0.002) (Figure 3c). Notably, by day 7 the difference was already clear. The AUC between day 2-7 showed significance in all SMX conditions (p < 0.01 for 1 μg/mL SMX, p < 0.002 for 4 μg/mL and 8 μg/mL SMX) (Figure 3d). The automated monitoring also collected speed data (Figures 3e,f), showing that the mean speed of moving worms was significantly slower for the SMX conditions at day 2.5 (p < 0.002, Supplementary Table S3) but speed across the whole experiment was significantly increased with SMX (Fig.f inset).

The WormGazer™ required disturbing the worms only three times over the time period while the manual lifespan required doing so every other day. No movement above the automated movement threshold was detected after day 22, whereas the manual method could detect smaller movements and response to prodding, and so lasted until day 40.

Overall, the two methods showed comparable results but with large differences in user effort and the amount of data captured. An AUC from days 2-7 showed the same significant result as a 40-day manual lifespan, which indicates that a 7-day healthspan can be used to detect lifespan extensions without requiring the work and time of a lifespan assay.

### Testing Alpha-Ketoglutarate

Next we tested alpha-ketoglutarate (αKG), a metabolic intermediate in the Krebs cycle shown to extend lifespan in *C. elegans* (Chin et al. 2014). To understand the effect of αKG on ageing we conducted a concentration-dependent effect on healthspan from 0.5 to 8 mM, with 8 mM being the reported effective concentration. We found that 8 mM in fact significantly reduced worm speed while 0.5 mM and 2 mM had no effect (Figure 4a).

**Figure 4.**
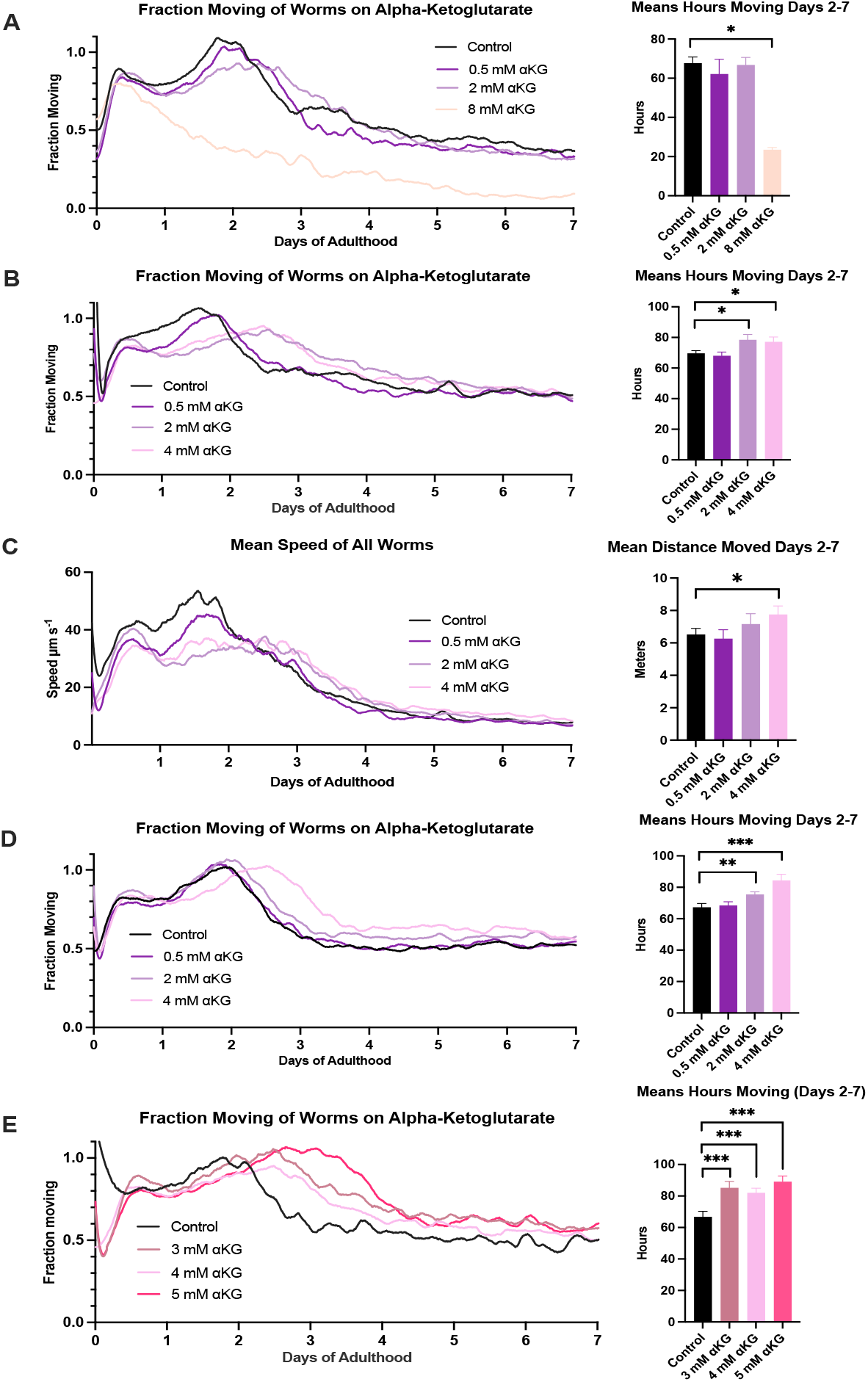
Alpha-ketoglutarate Improves Worm Health. (A) Fraction moving of worms on alpha-ketoglutarate and its integration of the area under curve (inset). n ≥ 180 worms, 6-8 petri dishes per condition. (B) Fraction moving of a smaller concentration range, identifying two positive concentrations with integration of AUC (inset). n ≥ 330 worms, 11-12 petri dishes per condition. Mean speed of all worms for the same experiment and integration of AUC (inset) (D) Fraction moving of a repeat, showing the same results with integration of AUC (inset). n ≥ 300 worms, 10-11 petri dishes per condition. (E) Fraction moving of a smaller concentration range, identifying three positively effecting concentrations with integration of AUC (inset). n ≥ 150 worms, 5-9 petri dishes per condition. * = p < 0.05, ** = p < 0.01, *** = p < 0.002, one-tailed test. Compound added to agar. Glp-4 worms used. Plotted in GraphPad.

We tightened the concentration range to 0.5 mM to 4 mM and used higher animal numbers, to find that 2mM and 4 mM αKG significantly improved movement by 12.7% and 10.7% respectively in the first 7 days compared to control (p < 0.002) (Figure 4b). Furthermore, 4 mM also improved mean speed of all worms (Figure 4c). The mean speed of moving worms for all experiments can be found in Supplementary Figure S2. This finding was reproducible (Figure 4d). An even smaller concentration range was then assayed to find that 3mM and 5 mM αKG were also effective in improving movement (Figure 4e).

By using a 7-day automated monitoring experiment, we found αKG is effective in maintaining health in a small range of concentrations.

## Discussion

### Movement Not Lifespan to Measure Ageing

In this paper we have presented a way to measure ageing in *C. elegans* using movement data rather than lifespan. This approach is made possible by the novel WormGazer™ automated imaging technology presented here, but other technologies that monitor movement across many worms at once non-invasively can be used to achieve a similar result. Manual lifespan assays have been used for decades because of the relative ease and transferability of the assay. It does not require any extra equipment than is found in a standard *C. elegans* lab, but only the skill and experience to handle the worms gently and to consistently score death.

The ubiquity of the assay has led to lifespan to be referred to almost synonymously with ageing. However, *C. elegans* lose optimal health within days of beginning their adult life, and thus ageing occurs long before death. The last week of a worm’s life is spent hardly moving at all and not eating (Newell Stamper et al. 2018). Maintaining health and/or preventing chronic disease is the goal of much research into ageing as well as drug development in the field, and thus an assay that addresses these endpoints directly is important.

### Validation and Reproducibility

In this paper we validated the approach of using movement to measure ageing by showing that the long-lived *age-1* mutant maintains movement with age longer than the wild type as measured by our assay. Simultaneously, we revealed features of early life movement of this mutant that are consistent with the idea that there are tradeoffs in IIS mutants that increase lifespan (Walker and Lithgow 2003).

Any method that measures ageing must critically be reproducible. Many environmental and epigenetic factors can affect health so it is critical to have very consistent culture conditions and worm preparation. Because the WormGazer™ monitors movement from the beginning of the experiment any anomalies in the worm behaviour can be detected before age-related decline begins, assisting quality control of the assay preparation. We have shown that the results from the WormGazer™ have great reproducibility (Figures 2 and 4, Supplementary Figure S1).

### Utility in Drug Development to Slow Ageing

Using the WormGazer™ technology it took only 7 days to see the age-slowing effect of SMX. We saw significant improvements with αKG after narrowing down from a wide concentration range. This finding illustrates the point that several concentrations of a compound should be tested to present the full picture and that automated imaging over 7 days makes dose response studies much faster and easier than using lifespan assays. The scalability of the technology coupled with an automated workflow to produce the petri dishes and worms in a standardised way, allows large numbers of new compounds to be screened at multiple concentrations or in combinations to find new interventions that slow ageing and improve human health. Once compounds that slow ageing have been identified, the power of *C. elegans* genetics can then be used with the same technology to uncover mechanism of action. In the process of drug development to slow ageing, safety is a very important factor. Any drug that is given to healthy people over the long term must be very safe, and monitoring movement throughout early to mid-adulthood highlights any potential early life toxicity. Overall we show that monitoring the movement of large populations of worms with time is a powerful tool to address the challenges of discovering new drugs that slow ageing.

## Materials and Methods

### Worm Maintenance

All strains are from the Caenorhabditis Genome Center (CGC) and maintained on agar Petri dishes with OP50 *E. coli* at 16 °C. The strains used were N2, TJ1052 (*age-1(hx546)* II), CF1038 (*daf-16(mu86)* I), and SS104 (*glp-4(bn2) II*).

To prepare for experiments on the WormGazer™, gravid worms were placed onto 9 cm petri dishes to lay eggs 4 days before being placed on the machine. The mothers are removed after 48 hours, and 24 hours later these 9 cm petri dishes are shifted to 24°C to ensure large numbers of L4s or to induce sterility in the case of SS104 (*glp-4(bn2)* worms that were sterile if raised at 20°C or above. On the following day, worms at the L4 stage were selected and 30 picked onto each 6 cm petri dish. At least 6 petri dishes were prepared per condition. The edges of these dishes were covered with parafilm to prevent desiccation, but a 5 mm gap was left for air exchange. Plates were then loaded into the machines and left undisturbed for the duration of the experiment.

### Preparation of Petri dishes

Petri dishes are filled with Defined Media (DM) in which peptone found in the standard Nematode Growth Medium (Brenner 1974) is replaced with defined amino acids and trace metals to minimise batch-to-batch variation in peptone as previously described (Virk et al. 2016) with vitamin B12 added (Maynard et al. 2018). Petri dishes for imaging are poured with DM agar 3 days before the worms are added. Floxuridine (FUdR) is added at final plate concentration of 2 μM to prevent progeny production in all strains that are not genetically temperature-sensitive sterile. All other compounds are made up in a 100x stock in required solvent and added to the agar before plates are poured. On the next day, petri dishes are seeded with 100 μl of an overnight culture of OP50 in LB. Petri dishes are stored at 20°C with controlled humidity until 24 hours after bacterial seeding, when they were transferred to an incubator at 24 °C.

### Imaging on the WormGazer™

For each camera, images are taken every 0.8s for a period of 160s to create a group of 200 images and stored on a single board computer. These images are converted into a series of analytical images on the single-board computer and those images are transfer to a central server. After 5 minutes, the process is repeated. From these analytical images, the number of moving objects is calculated by applying a threshold of the minimum speed of each object of 10 μm/s. The speed is derived from the length of the object divided by the 160 second time interval of the imaging window.

### Manual Lifespan Protocol

SS104 *glp-4(bn2)* worms are prepared in exactly the same way as for healthspan experiments. L4 worms are selected and picked onto petri dishes with 30 worms per dish and 12 dishes per condition. Worms are scored every second day during weekdays from day 7 onward, and marked as alive, dead, or censored. Worms are prodded with a platinum pick to check for movement before being marked as dead. Worms are transferred on days 7 and 14. Censored indicated the worms had bagged, burst vulvas, or crawled off the plate. A Kaplan-Meier curve was generated on GraphPad Prism and statistics on JMP statistical software.

### Statistical Analysis

The shading on the time-series curves is one standard error of the mean across the curves for each petri dish within the condition. The error bars on the bar charts are ± 1 standard error on the mean across the measurements for each dish within the condition. When comparing two conditions on a bar chart, the difference between the means is calculated, and the standard error on this is calculated with reference to the standard error on each individual measure, using gaussian error statistics. The significance thresholds are then set with reference to the difference expressed in terms of its standard error. A difference of less than 1.64 standard errors is marked as not significant (ns). A difference between 1.64 and 2.33 standard errors is marked as one star (*), corresponding to P<0.05 on a one-sided test. A difference between 2.33 and 2.83 standard errors is marked as two stars (**), corresponding to P<0.01 on a one-sided test. A difference greater than 2.83 standard errors is marked as three stars (***) corresponding to P<0.002 on a one-sided test.

## Acknowledgements

We thank Michael Fasseas, Alana Mullins, Sushmita Maitra who assisted with technical aspects of the experiments, Craig Manning who built and contributed to the design of early prototypes of the technology, Fozia Saleem for providing useful feedback on the manuscript.

## Author contributions

GZ, DW conceived of study, designed C. elegans experiments and wrote the manuscript. CS conceived of, designed and built the WormGazer technology. AK helped with design and building of the technology. AR, GZ helped design and executed the C. elegans experiments.

