## Supplementary Data for "Measuring *C. elegans* Ageing Through Non-Invasive Monitoring of Movement across Large Populations"

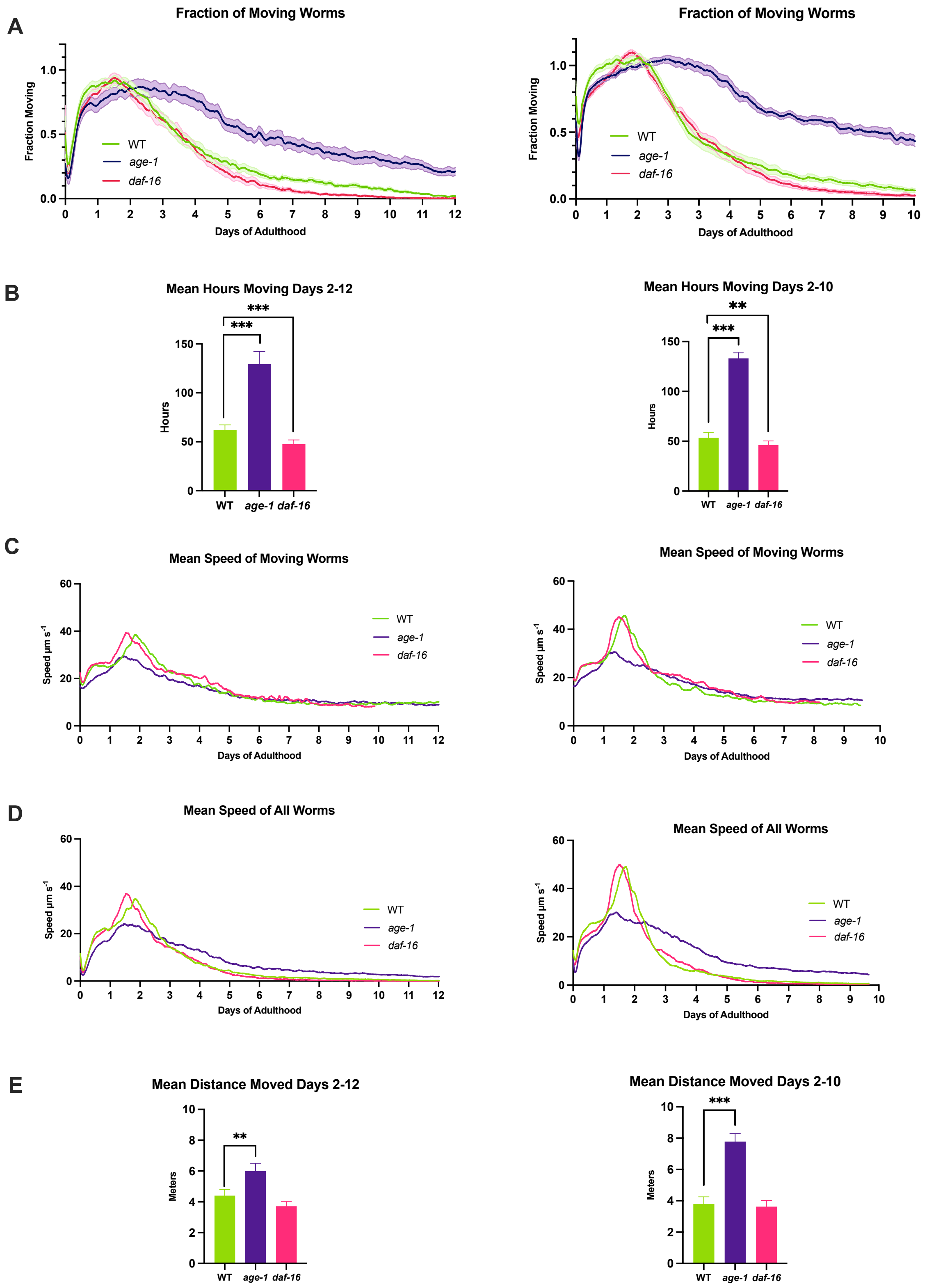


*Supplementary Figure S1 Direct Comparison of Two Repeats of age-1, daf-16 and WT worms. (A) The fraction moving graph shows the proportion of worms moving during the imaging window. (B) The area under the curve integration from day 2 to end of experiment for (A). (C) The mean speed of moving worms. (D) The mean speed of all worms, which is a function of A and E. (E) The area under the curve integration from day 2 to end of experiment for (D). Lefthand column; n ≥ 298 worms, 10-12 petri dishes per condition. Righthand column; n ≥ 420 worms, 14 petri dishes per condition. Plates with 2 µM FUdR on DM. *** = p <0.002, one-tailed test. Plotted in GraphPad.*

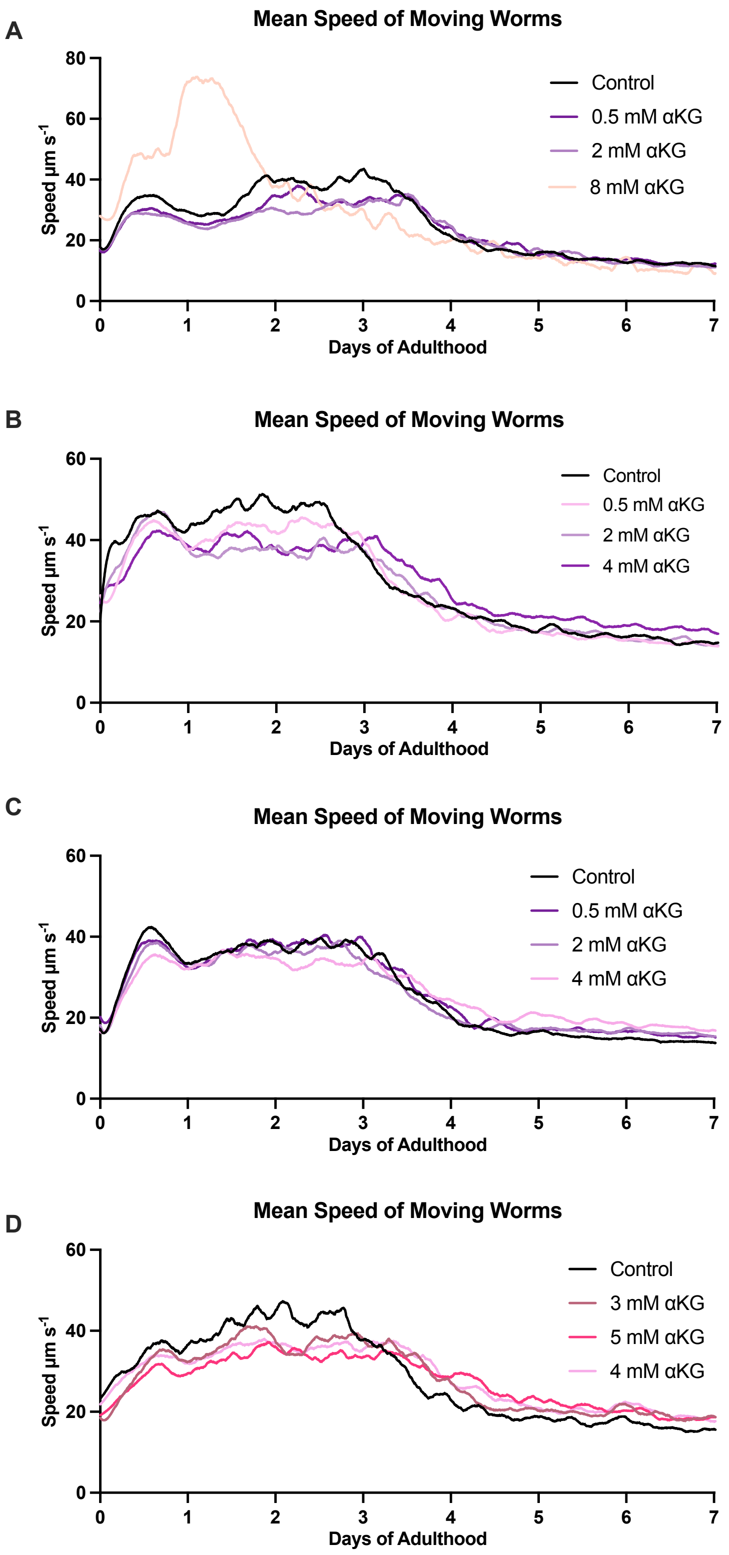


*Supplementary Figure S2 Mean Speed Moving Worms for Alpha-ketoglutarate. Graphs are in order of appearance in Fig. 4. (A) Mean speed of moving worms. n ≥ 180 worms, 6-8 petri dishes per condition. (B) Mean speed of moving worms. n ≥ 330 worms, 11-12 petri dishes per condition. (C) Mean speed of all worms. n ≥ 300 worms, 10-11 petri dishes per condition. (D) Mean speed of moving worms. n ≥ 150 worms, 5-9 petri dishes per condition. Compound added to agar. Glp-4 worms used. Plotted in GraphPad.*

*Supplementary Table S1 Manual lifespans were carried out at 24˚C. Worms were checked every other day on weekdays after day 7. SS104 glp-4(bn2) worms used, with OP50 E. coli on DM plates. Work carried out with assistance from Adelaide Raimundo. Statistics by JMP. All comparisons against control.*

| Condition | n | Censor | Mean Lifespan | Standard Error | % Change | p-value (Log-Rank) | p-value (Wilcoxon) |
|---|---|---|---|---|---|---|---|
| Control | 290 | 100 | 18.39 | 0.28 | - | - | - |
| 1 µg/mL SMX | 276 | 107 | 20.28 | 0.28 | +10.27% | < 0.0001 | < 0.0001 |
| 4 µg/mL SMX | 276 | 108 | 20.99 | 0.36 | +14.13% | < 0.0001 | < 0.0001 |
| 8 µg/mL SMX | 262 | 124 | 21.14 | 0.36 | +15.00% | < 0.0001 | < 0.0001 |

*Supplementary Table S2 Raw Data for the Percentage Comparisons for age-1, daf-16 and WT. Data taken from machine export files.*

| Condition | Fraction Moving AUC Days 0.5-2 | Mean Speed Moving Day 2 |
|---|---|---|
| Control | 31.62 ±1.30 | 0.89 ± 0.05 |
| *age-1(hx546)* | 27.78 ± 1.87 | 0.85 ± 0.06 |
| *daf-16(mu86)* | 30.48 ± 1.34 | 0.86 ± 0.04 |

*Supplementary Table S3 S4 Raw Data for the Percentage Comparisons of Mean Speed of Moving Worms in Automated Lifespan. Data taken from machine export files.*

| Condition | Mean Speed Moving Day 2.5 |
|---|---|
| Control | 49.68 ± 2.12 |
| 1 µg/mL SMX | 40.39 ± 1.46 |
| 4 µg/mL SMX | 34.37 ± 1.39 |
| 8 µg/mL SMX | 32.48 ± 1.37 |
